# Microbial diversity ensures the stability of carbon cycling processes under increasing temperature in model soils

**DOI:** 10.1101/2022.12.19.521036

**Authors:** Nadja Rohner, Selina Lepori, Viviana Loaiza, David Sebag, Eric Verrechia, Daniel B. Nelson, Ansgar Kahmen, Pascal A. Niklaus, Anna-Liisa Laine, Luiz A. Domeignoz-Horta

## Abstract

Biodiversity loss and climate change are two of the most alarming threats to ecosystem functioning and stability. These factors are, however, typically investigated separately, overlooking potential interactions between species extinctions and climate change on ecosystems. Here, we evaluated how different temperature regimes impact the relationship between microbial diversity and ecosystem functioning, in terms of temperature sensitivity of carbon (C) cycling functions. We hypothesized that more diverse soil communities promote the stability of C cycling functions under both temperature regimes. We did not observe a ubiquitous response of all C-cycling processes to temperature increases within the distinct community diversities. While growth was stable with increasing temperatures among the diversity levels, respiration rates increased more strongly at low diversity levels than high diversity levels at higher temperatures. Carbon use efficiency, which integrates both growth and respiration, tended to decrease with temperature at lower levels of diversity. Jointly, these results demonstrate the importance of belowground soil community diversity for maintaining C-cycling thermal response under changing climate.

## Introduction

To date, most studies on the ecological consequences of biodiversity loss have focused on above-ground processes, although a large part of terrestrial biodiversity is hidden within the soil (Wagg *et al*., 2014). While soil microorganisms are the most diverse and most abundant organisms on the planet and are crucial drivers of a wide range of ecosystem functions (Singh *et al*., 2010), the consequences of belowground microbial diversity loss for belowground ecosystem functions are not well understood. This is critical given the numerous anthropogenic threats to soil biodiversity (Wagg *et al*., 2014, Cavicchioli *et al*., 2019, Gallego *et al*., 2021, Gallego and Martin-Laurent, 2020, Zak *et al*., 2003). A belowground ecosystem function that is increasingly more studied due to its relevance for the climate system is soil carbon (C) cycling (Cavicchioli *et al*., 2019, Domeignoz-Horta *et al*., 2022, Cappelli *et al*., 2022). Empirical data provided by Kallenbach *et al*. (2016) suggests that soil microbial physiology may be more important for soil carbon stabilization than traditionally assumed. Accordingly, it has been recently shown that community growth efficiency was positively related to bacterial diversity under favorable environmental conditions (Domeignoz-Horta *et al*., 2020). Further, it was also shown that distinct microbial communities generate soil organic matter (SOM) with different chemistry (Domeignoz-Horta *et al*., 2021), suggesting that even broad C-cycling-related processes in soil might be shaped by microbial diversity and composition contrarily to what has been previously acknowledged (Schimel *et al*., 2012).

Growth efficiency or carbon use efficiency (CUE) is an important parameter to characterize microbial physiology (Manzoni *et al*., 2012, 2018) and determines the partitioning of plant inputs into a fraction of carbon remaining in the soil (microbial growth) and a fraction respired as carbon dioxide to the atmosphere. The CUE of microbial communities does not only depend on the identity of an organism but also on the interactions with other community members, and therefore on community composition and diversity (Domeignoz-Horta *et al*., 2020). CUE is also affected by abiotic factors such as temperature, moisture and nitrogen availability (Timmis *et al*., 2019). A seminal study showed that changes in temperature systematically modified the relationship between diversity and community growth (García *et al*., 2018). Moreover, a recent study showed that the increase in respiration in response to warmer temperatures was higher in low diverse soils (Xu *et al*., 2021). These new findings suggest that more species are needed to maintain ecosystem functioning and stability under thermal stress. How all these factors interact and influence community CUE still remains largely unclear despite its importance for understanding soilatmosphere carbon cycling feedback under global change (Allison *et al*., 2010).

## Materials and Methods

Here, we experimentally test the hypothesis that more diverse microbial communities temporally stabilize CUE and the underlying processes growth and respiration when exposed to temperature variation (Loreau *et al*., 2001). We used a sterile, carbon-free model soil as it has been shown that such a soil can provide a platform to isolate specific components from confounding factors compared to natural soils (Kallenbach *et al*., 2016). The model soil is composed of sand and clay (9:1) and was prepared as described by Domeignoz-Horta *et al*. (2020). Soil microcosms consisted of 50 ml falcon tubes filled with fifteen grams of this substrate and inoculated with 3 ml microbe-media (Tryptic soy broth; TSB) solution, reaching 70% of the water holding capacity. We created a microbial diversity gradient by serially diluting a community obtained by extracting an agricultural soil (D0; no dilution > D1; first dilution > D2; second dilution) (Figure 1, Supplementary material) (Diaz *et al*., 2003). The least diverse treatment (D3) is represented by a single bacterial strain commonly found in soils (*Streptomyces* sp. DSM 687; Supplementary material).

**Figure 1.**
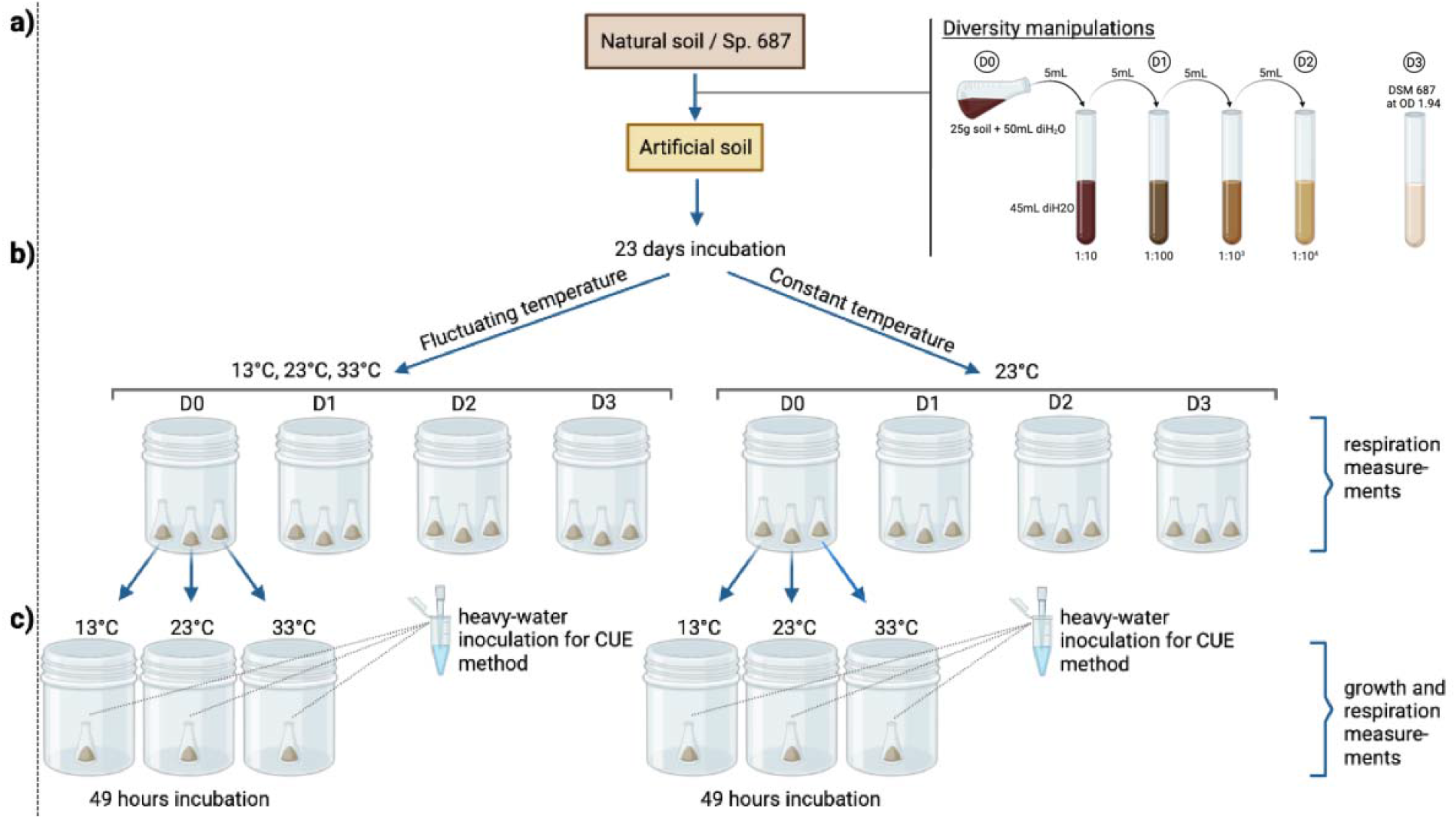
Experimental design. The microbial diversity manipulation of the soil suspension was performed to obtain D0, D1 and D2 inocula, while *Streptomyces* sp. DSM 687 was used for D3 treatment **(a)**. The microbial communities obtained with the diversity manipulations were inoculated into a model soil with 5x TSB media and incubated for 23 days under two temperature regimes **(b)**. Growth and respiration were measured after 49 hours of incubation at all temperatures **(c)**.

The experiment consisted of two phases, community establishment (phase I) and C-cycling evaluation (phase II). During phase I of the experiment, Figure 1b), constant (23°C) and fluctuating temperatures (increasing in temperature every 48 h: 13° -> 23° -> 33°C and restarting the cycle at 13°C) were applied to induce changes in the metabolism of microorganisms and select for microbial communities with distinct thermal niches (Schnyder *et al*., 2020). In phase I, we used four biological replicates for each diversity level at each temperature regime (4 biological replicates x 4 diversity treatments x 2 temperature regimes = 32 samples) and measured respiration as a proxy for microbial activity (Supplementary Material). At the end of phase I, we added fresh substrate (1 mg C per g soil with a solution consisting of TSB and glucose) to ensure that substrate was not limiting microbial growth. In the following phase II, we measured growth and respiration (Figure 1c) at 13°, 23° and 33°C in all soils (4 biological replicates x 4 diversity treatments x 2 temperature regimes x 3 incubation temperatures= 96 biological samples). At the end of the 48 h incubation, we measured respiration by headspace sampling with a gas chromatograph (Agilent Technologies 7890A) and growth in a subset of 0.9 g of soil with the ^18^O-H_2_O water method (Supplementary Material) (Spohn *et al*., 2016). Duplicate DNA extractions for each biological replicate were performed and CUE was calculated as by Spohn *et al*., (2016) (Supplementary material).

## Results & Discussion

### Microbial activity during community establishment (phase I)

The microbial respiration (CO2-C) decreased over the course of the 23-day incubation, in both the constant (CT) and the fluctuating (FT) temperature treatments. In CT, respiration decreased steadily over time except in the D2 treatment in which respiration increased towards the end of the incubation (last two measurements) (Figure 2A-B). This effect is likely due to the higher dilution and therefore smaller community inoculated into the soil which resulted in more activity towards the end of phase I, as substrate and niche space were not limited in comparison to less diluted treatments. In FT respiration clearly shows a thermal-response to incubation temperature (e.g. increasing when temperature increases and decreasing when the temperature decreases; Figure 2B). The diversity effect on cumulative respiration is larger in the CT (D0 & D1 > D2 & D3) than in the FL treatment, where no clear pattern is visible (Figure 2C).

**Figure 2:**
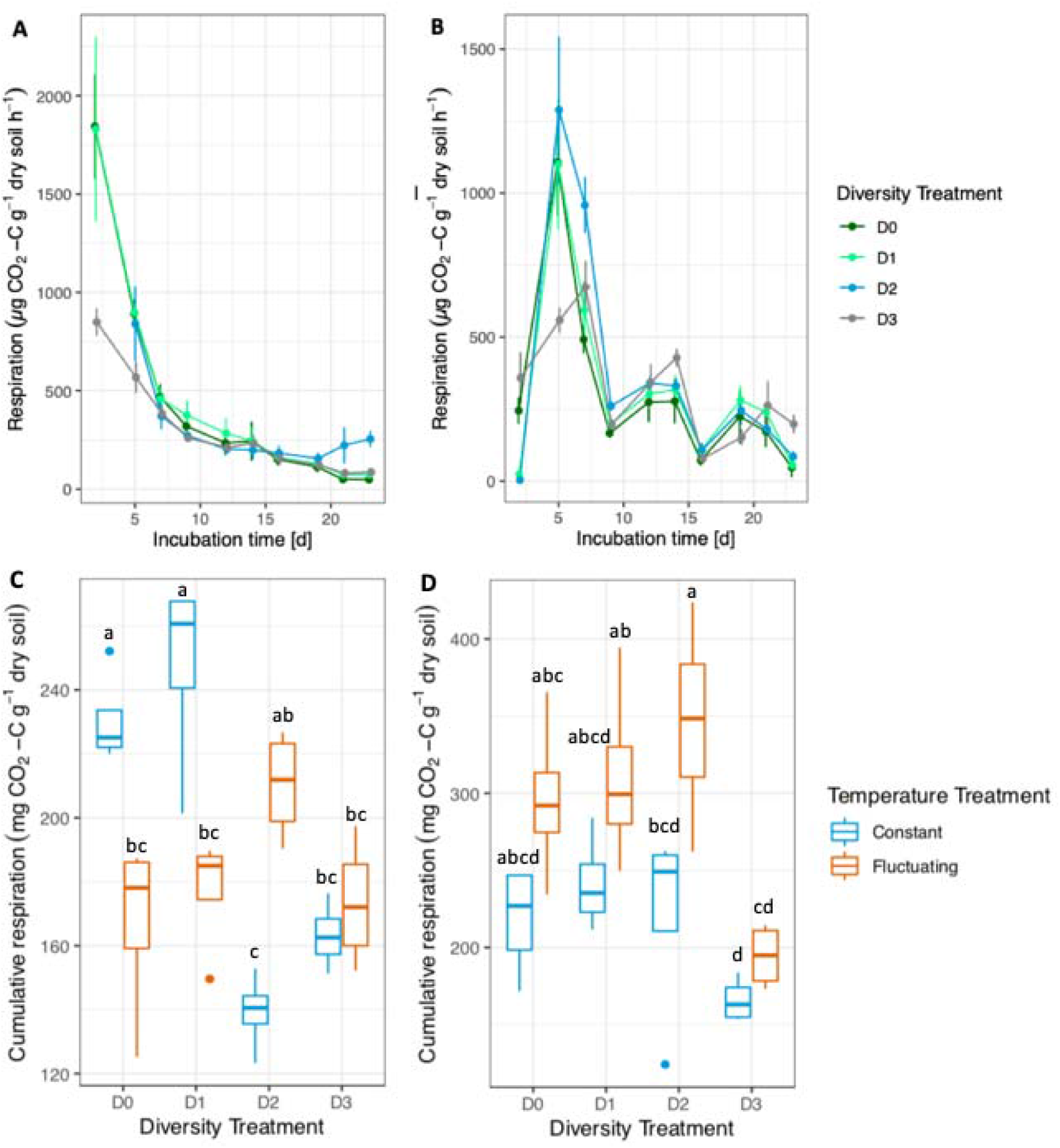
Microbial respiration rates during community establishment. Soil respiration of microcosms incubated at constant temperature (23°C) **(A)**, or fluctuating temperatures (13°C, 23°C, 33°C) **(B)**. For A and B error bars denote standard deviations of biological replicates (n = 4). Cumulative respiration for the overall incubation period **(C)** and cumulative respiration for the time-points chosen when the fluctuating temperature treatment was incubated at 23°C **(D)** allowing comparison of respiration rates at the same temperature between both treatments. For boxplots, the box denotes interquartile range, whiskers denote the 1.5 × interquartile range. The horizontal line denotes the median. Points represent outliers outside of the 1.5x interquartile range. Letters indicate significant differences between diversity levels tested with the Tukey-HSD test.

In addition to the total cumulative respiration, we also calculated the cumulative respiration for the time points in which microcosms of the CT and FT treatments were both at 23°C (Figure 2D). We observed a difference in the cumulative respiration by both diversity treatment (F = 8.13, *p* < 0.001) and by temperature treatment (F = 18.23, *p* < 0.001) (Figure 2D) showing that the temperature regime influenced microbial activity also when evaluating the activity at the same common temperature. One possible explanation is that the 10°C temperature shift of the FT soils (as they were incubated at 13°C before 23°C) induced a stronger microbial respiration response in FT compared to CT (Figure 2D) as respiration is expected to respond exponentially to increases in temperature (Alster *et al*., 2020). The FT regime might have also resulted in a larger community size and consequently showed higher activity compared to the other treatment. Previous studies with model soils have shown that abiotic factors drive the community composition (Kallenbach *et al*., 2016, Domeignoz-Horta *et al*., 2020). Thus, another potential explanation is that the distinct temperature regimes selected communities with different thermal niches, and therefore these communities show distinct respiration rates at the same temperature (Figure 2D). Accordingly, Pold *et al*. (2020) showed that the response of soil isolates to temperature is strain specific varying with temperature and substrate.

### Soil C-cycling response to different temperatures and diversity levels (phase II)

After the microbial communities were exposed to distinct temperature regimes, in Phase II we incubated them under 3 different temperatures and measured CUE. We used linear mixed models to evaluate how the different diversity levels and the temperature regimes impacted the different soil C-cycling functions (Figure 3). Growth did not depend on temperature in both temperature regimes and diversity levels (Figure 3A and D). Respiration, however, increased faster with increasing temperatures at low levels of diversity compared to higher diversity levels (Figure 3B and E) which has also been reported in natural soils (Xu *et al*., 2021). This pattern was consistently independent of previous temperature regimes.

**Figure 3:**
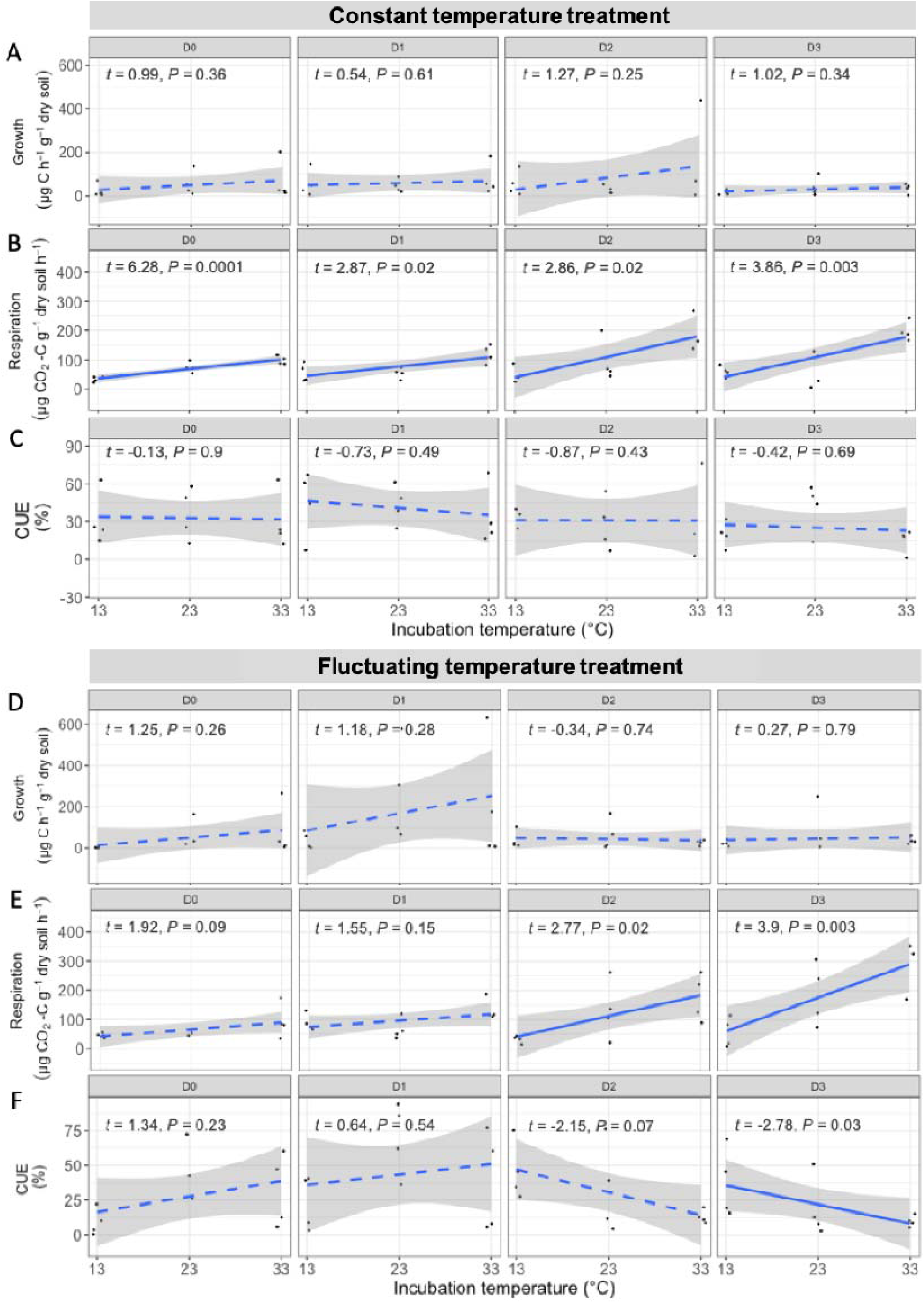
Relationship between C-cycling processes and temperature within the different diversity levels for the constant (A, B and C) and fluctuating (D, E and F) temperature regimes. Linear mixed effect models were used to evaluate the impact of short-term temperature changes on respiration, growth and CUE. Linear curves were fitted when the result is significant or dashed line if not significant.

We hypothesize that the serial dilution method could have affected the strategies of microbial communities in the sense that higher dilutions, paired with readily available nutrients lead to a community more strongly dominated by copiotroph-strategists and, therefore a higher respiration response to temperature. While copiotrophs-related strategists have been described to be fast-growing and have a low carbon use efficiency, oligotrophs-related strategists are slow-growing and have a high carbon use efficiency (Fierer *et al*., 2007; Leff *et al*., 2015). The higher diluted soils likely have a low diversity of copiotrophic-species while in the less diluted we have a higher species richness which will interact with each other (e.g. compete for the resources) allowing a broader spectrum of copiotroph-related physiologies and distinct strategies to emerge (Song *et al*., 2017). Accordingly, Bai *et al*. (2017) suggested that respiration response to a 10°C increase in temperature (Q_10_) could be controlled by the shift between copiotrophs-like and microbial strategies. Regarding the D3 treatment, where the soil was inoculated with a single *Streptomyces* strain, very high respiration Q_10_ values were observed (Fig. 3 B, E). This species belongs to the phylum Actinobacteria which exhibits copiotrophic attributes (Fierer *et al*., 2007).

It is often considered that high diversity is required to ensure ecosystem functioning (Garcia *et al*., 2018, Maron *et al*., 2018, Yang *et al*., 2021). Moreover, García *et al*. (2018) show that high diversity is needed to ensure growth in more extreme temperatures and that high diversity maintains high growth potential. These results were obtained from marine microorganisms under liquid cultivation, while our experiment using a model soil introduces some heterogeneity to simulate the natural gradients observed in soils (e.g. substrate, oxygen, diffusion rates). In contrast to these previous results, we show that absolute growth and growth Q_10_ was not different among diversity treatments (Figure 3A, D). Growth efficiency (CUE) is the combination of growth and respiration, if respiration increases faster with temperature at low levels of diversity but growth remains constant, CUE is expected to decrease at low levels of diversity with increasing temperatures. We observed a trend of decreasing CUE with increasing temperatures and decreasing diversity (Figure 3C and F). However, due to high variability in the data this was only significant at D3 and marginally significant at D2 diversity levels within the FT treatment (Figure 3F). We measured CUE as a microbial physiology parameter that is considered important to understand the fraction of C that potentially stays within the soil in relation to what is respired (Allison *et al*., 2010). Our results show that the Q_10_ of respiration and growth responded differently along the diversity gradient resulting in a decreasing CUE with increasing temperature as previously reported (Frey *et al*., 2013). Additionally, our results suggest that the CUE Q_10_ decreased more in the less diverse communities, supporting a previous observation that a more diverse community was able to respond to changes in temperature and/or moisture without decreasing its growth efficiency (Domeignoz-Horta *et al*., 2020).

To further evaluate the stability of C-cycling functions between the different diversity levels across changes in temperatures, we evaluated the coefficient of variation compiling all C-cycling measurements (Figure 4) for both temperature regimes (FT and CT) as previously (Eisenhauer *et al*., 2011) and the 95% confidence interval (Supplementary Figure 1 and 2). We refer to the inverse of the coefficient of variation as the “C-cycling stability” to temperature variations. We observed no significant differences between the fluctuating and constant temperature regimes (Figure 4). The stability of C-cycling to temperature variations did not depend on temperature regime. However, C-cycling functions were less stable at low diversity, (Figure 4 and Supplementary Figure 1 and 2). Interestingly, compiling all our individual measurements within one diversity level leads us to support the functional redundancy hypothesis of C-cycling (Schimel *et al*., 2012). However, we observed a distinct process rate response to different temperatures among diversity levels for respiration. Considering that these contrasting results were obtained in a simple model soil system, it highlights the need to evaluate various ecosystem functions when evaluating the diversity x ecosystem function relationship in complex natural soils if we aim to have a deeper understanding on how C-cycling in soils may change following environmental and anthropogenic perturbations. Moreover, Yang *et al*. (2022) show that multiple anthropogenic pressures can modify communities and soil functioning, and highlights the importance of incorporating multi-factor experimental approaches considering that about 98% of studies still evaluate a single, or two factors only (Rilling *et al*., 2019). Here we used a 3 multi-factor experimental approach (diversity loss x temperature regime x temperature response of C-cycling functions) to unravel how microbial-controlled C-cycling functions may shift in response to changes in their environment. Altogether, our results highlight that successive changes (diversity loss associated with temperature increases) can result in changing ecosystem process rates of importance for soil C-cycling.

**Figure 4:**
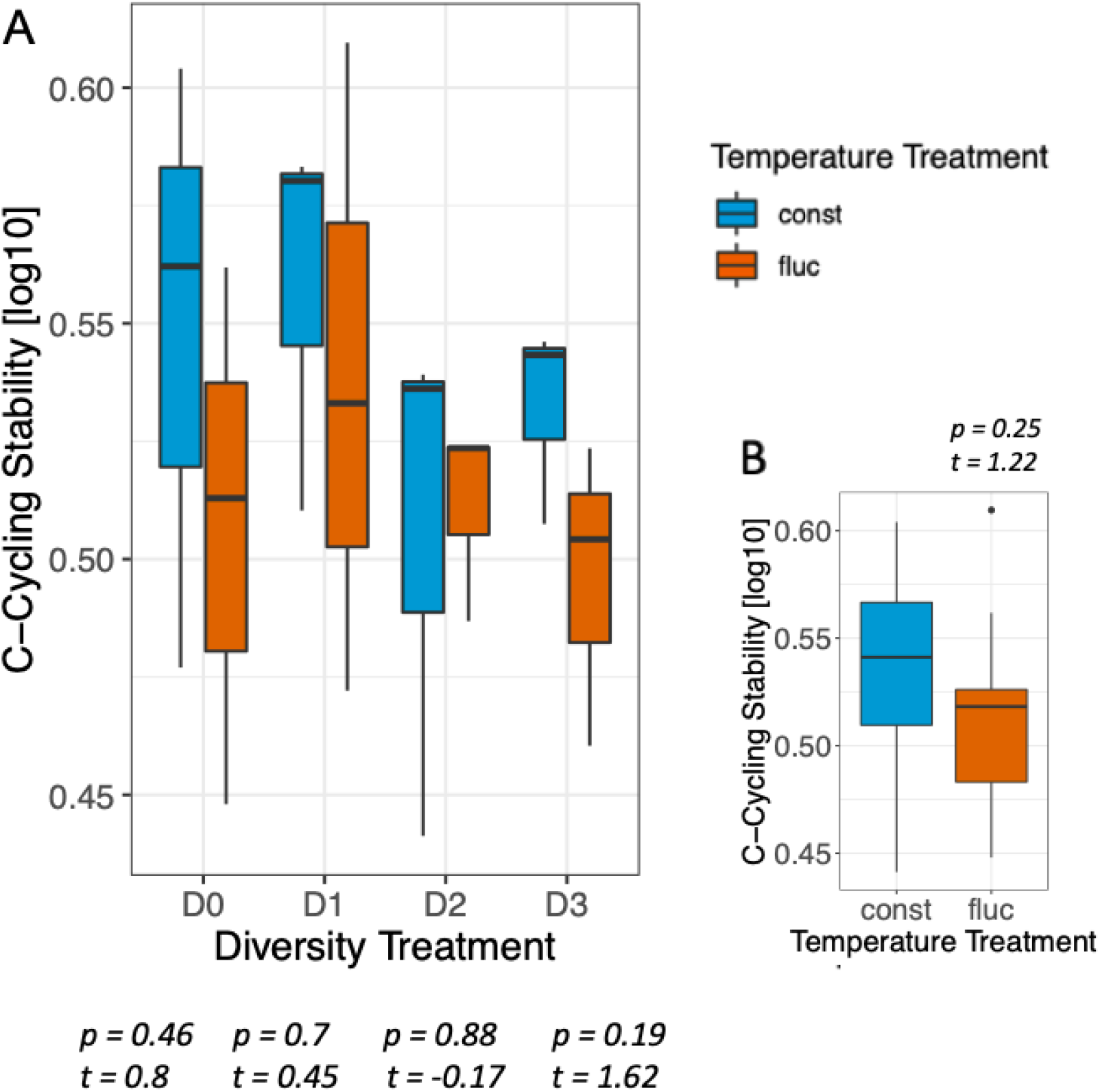
C-cycling stability to changes in temperature. Coefficient of variation compiling all C-cycling functions between diversity treatments and temperature regimes **(A)** and between the two temperature regimes **(B)**. The box denotes the interquartile range, whiskers denote the 1.5× interquartile range. The horizontal line denotes the median. Points indicate values outside of the 1.5x interquartile range. Letters indicate significant differences (*P* < 0.05) between diversity levels tested derived from the Tukey-HSD test.

## Acknowledgements

Funding for this project was provided by the Academy of Finland (STN Multa; 327222) to A-L.L. This work was also conducted with support from the University of Zurich. We would also like to thank the Laine Lab team members for the supportive discussions and Prof. Dr. Owen Petchey for being a great statistics teacher.

## Notes

### Competing Interest Statement

The authors have declared no competing interest.

