## Supplementary material for "Microbial diversity ensures the stability of carbon cycling processes under increasing temperature in model soils"

**Supplementary figures:**

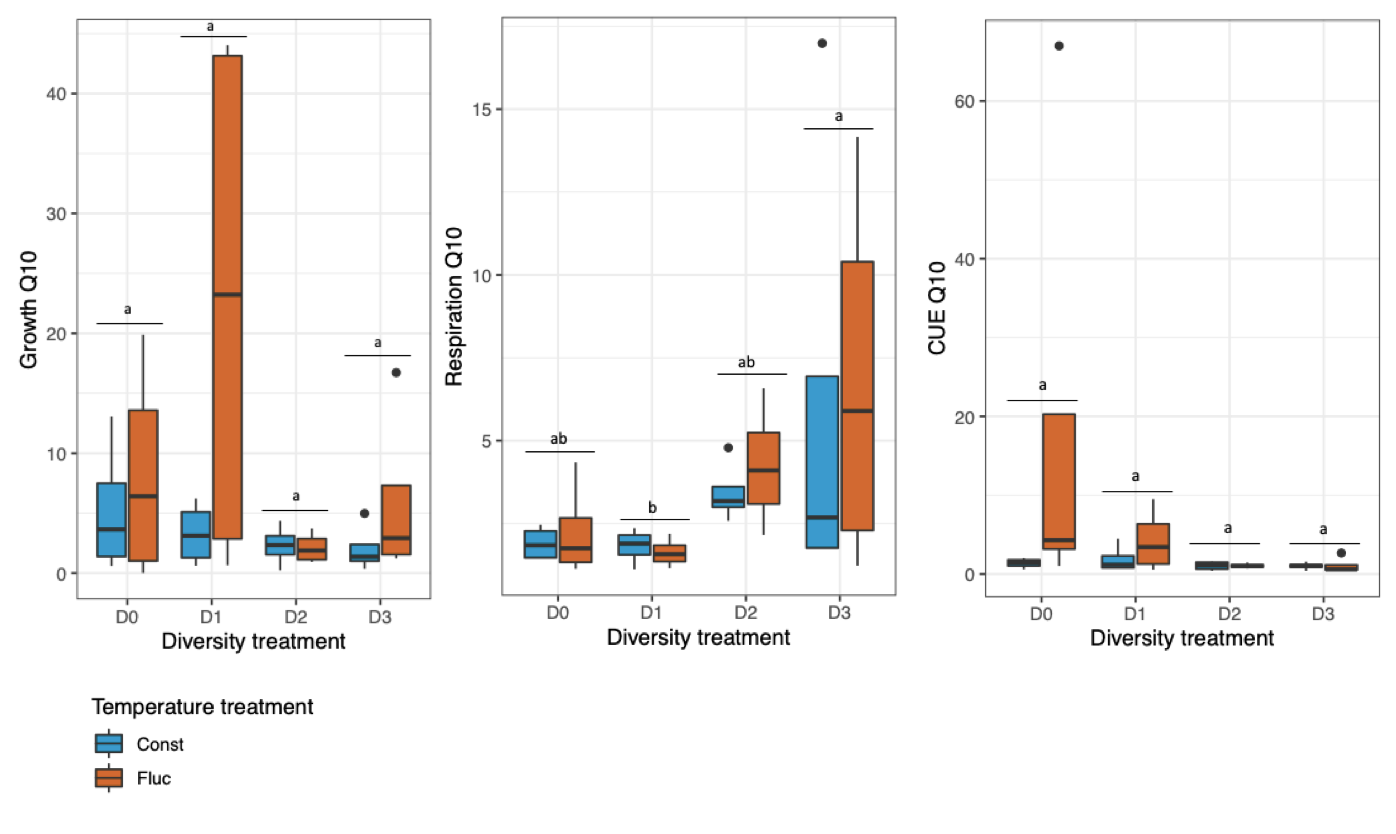

**Supplementary Figure 1. Response of C-cycling processes to a 10^o^C increase in temperature (Q10)**. Temperature sensitivity for the distinct C-cycling processes: Q_10_ of growth, respiration, and CUE at the two temperature regime treatments within all diversity levels. Anova followed by Tukey HSD test was used to detect significant differences between diversity levels regarding the different C-cycling processes.

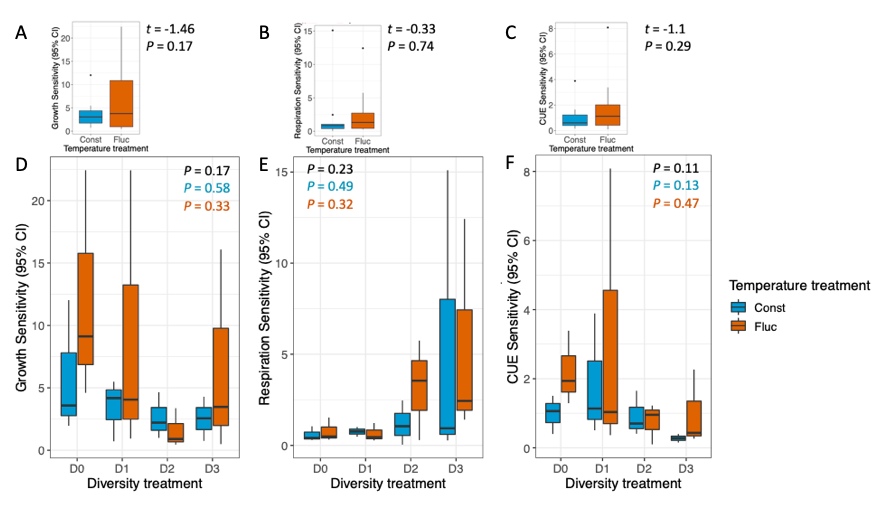

**Supplementary Figure 2. Variability of the temperature sensitivity of C-cycling processes**. 95% CI of the temperature sensitivity (Q_10_ value) for the distinct C-cycling processes. In the upper panel, 95% CI of Q_10_ of growth (A), respiration (B) and CUE (C) between the two temperature regime treatments within all diversity levels. A Welch Two Sample t-test was used to evaluate statistical differences. In the lower panel, 95% CI of Q10 of growth (D), respiration (E) and CUE (F) between diversity levels. Anova followed by Tukey HSD test was used to detect significant differences between diversity levels and the response variable was log transformed. The p-value compares the diversity treatments for constant and fluctuating conditions (black), constant conditions only (blue) and fluctuating conditions only (orange). For each boxplot, the box denotes the interquartile range, whiskers denote the 1.5× interquartile range.

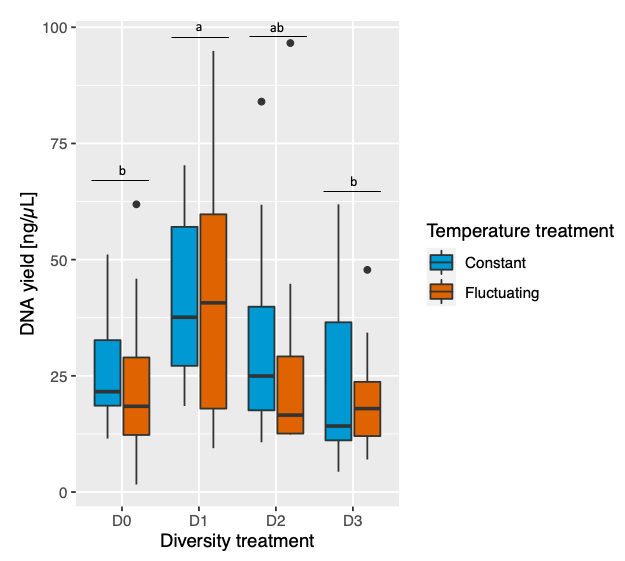

**Supplementary Figure 3. DNA yield at the end of incubations**. DNA concentration for the constant temperature treatment only (blue) and fluctuating temperature treatment only (orange). Anova followed by Tukey HSD test was used to detect significant differences between diversity levels and the response variable was log transformed. For each boxplot, the box denotes the interquartile range, whiskers denote the 1.5× interquartile range.

**Supplementary Text**

#### **Soil inoculation**

Each soil microcosm contained 15g (dry mass) artificial soil which was inoculated with a 3ml microbe-media blend in order to reach a WHC of 70%. The blend containing microbes (D0-D3) was mixed 1:1 of media and inoculum to target ~ 1mg C ^-1^ g soil per week of incubation. The microcosms were inoculated in the biosafety cabinet in order to avoid contamination and the flask opening was covered with a parafilm layer directly after inoculation. Thereafter they were stored at three samples each in a jar in the incubator with the respective temperature (Figure 1). Furthermore, six non-inoculated controls injected with 3ml TSB were set up to control for contamination. Respiration measurements were taken during the 23 days of incubation.

#### In preliminary experiments different media types were tested, including the media recommended by the German Collection of Microorganisms and Cell Cultures (DSMZ), where the bacterial strain was ordered. For the experiment the tryptic soy broth (TSB) was used. For the liquid media 30g TSB was mixed with 1000 ml distilled water and autoclaved in a wet-cycle. This liquid media was used to grow Streptomyces sp. (DSM 687) to a previously determined optical density.

#### **Strain Preparation**

The Streptomyces species (DSM 687) ordered at DSMZ was inoculated as a single species and represents the least diverse treatment. It arrived freeze dried as a small pellet in a glass vial. The pellet was soaked with 500μl of the recommended Gym Streptomyces Media and from that stage plated with a loop on agar plates and suspended several times in 100ml of liquid media. Optical density (OD) was measured at 600nm over the period of 5 days in order to obtain a growth curve. Freezer stocks were created at OD 1.6 which lies within the exponential growth phase.

In preliminary experiments we evaluated that the Streptomyces sp. (DSM 687) grows well in the model soil under all the temperatures. Its preferred temperature in liquids lies at 28°C. Regarding dominant bacteria in natural soils, Delgado-Baquerizo et al. (2018) conducted a global analysis of bacterial communities in surface soils, where they identified the dominant bacterial phylotypes from 237 locations. They identified that the phyla Actinobacteria and Acidobacteria are one of the most abundant bacteria occurring in these soils. Furthermore, Streptomyces sp. which belong to the phylum Actinobacteria are important regarding carbon cycling (Barka et al., 2016). Regarding the more diverse microcosms, the microbes were extracted from a natural agricultural soil. Serial dilutions were made in order to create different diversity as previously (Diaz et al., 2003).

**Respiration measurements**

Soil respiration was used as an indicator for microbial activity and measured by gas chromatography (GC). For the respiration measurement the microcosms were placed in air-tight glass jars (volume = 900mL) with lids equipped with a valve allowing gas sampling with a gauge needle attached to a 60mL syringe. All jars were distributed randomly on a table and ventilated for several minutes in order for the CO_2_ concentration to be the same in all jars. Two room controls were closed at the same time in order to determine the ambient air and subtract that initial value from all the measurements taken at that time. All jars were closed airtight within two minutes and put back to the respective incubator. One and two hours after closing the jars, air samples were extracted. For this purpose, the jars were brought back to room temperature for approximately 10 minutes. For each sample 25mL air was extracted from the headspace of the jar after flushing the syringe 2 times at 5mL to mix the air in the jar. The 25mL sample was then released into a 12mL vacuum-sealed glass vial. Finally, the jars were opened to prevent the air to get anoxic and put back into the respective incubator. After the sampling period the glass vials were analyzed for their CO_2_ concentration in ppm with a gas chromatograph (Agilent Technologies 7890A) fitted with an automatic sample handler (Gilson GX-271). The samples were calibrated to four CO_2_ standards of 1050.1, 734.2, 408.4 and 18.2 ppm. A linear regression line was established for each measurement (evaluated from three points being t0 = room sample which is a control jar closed when closing the other samples, t1 = measurement after 1h, t2 = measurement after 2hrs) in order to detect any variability that was due to measurement errors of the gas chromatograph. All measurement points with an R^2^ value greater than 0.9 were retained which resulted in a removal of 16 measurement points. That threshold was chosen, as many points were in the range of an R^2^ value between 0.9-1. The values of the deleted points were much lower than those 90% and would have confounded the data.

#### **Addition of 18O-water for CUE measurements**

One day prior to 18O water addition, 0.6g of soil was collected from the 15g microcosm, weighted and put into microfuge tubes. The tubes were left open overnight in the biosafety cabinet allowing some water to evaporate. The weight was measured the next day and ^18^O-H_2_O diluted with unlabelled de-ionized water was added to reach 20% of final water content. The two microcosms (0.6g and 14.4g) were stored together in one jar for the respiration measurements. This decision was taken as the 0.6g microcosm might have yielded in CO_2_ values too small to measure. The jars were closed air-tight immediately after water addition and put into the respective incubator.

#### **DNA Extraction**

Genomic DNA was extracted from 0.6g soil using the QIAGEN DNeasy PowerSoil HTP 96 Kit (384) following the protocol of the manufacturer in duplicate extractions. The resulting DNA concentration was quantified fluorometrically using ThermoFisher Scientific, Quibit 4 Fluorometer. The DNA samples were dried in a silver capsule with a 7.5ml salmon sperm DNA solution and sent to the stable isotope ecology lab in Basel that analyzed the amount of δ^18^O in the samples.

### **Respiration and CUE calculations**

The respiration rate [ppm] was converted to CO_2_-C equivalent for each jar. From the measured CO_2_ in the headspace, a room control was subtracted. For conversion from ppm to mL the amount of substance n was derived from the ideal gas law:

$$pV=nRT$$

(2)

Where p is the pressure [atm], V is the volume [mL] of the measurement taken from the headspace, n is the amount of substance in the gas, R is the ideal gas constant which equals 82.06 and T is the temperature [K].

In a next step the CO_2_-C_Equivalent_ [ml g^-1^ h^-1^] was calculated by the following equation

$${CO}_{2}{-C}_{Equivalent}= \frac{{CO}_{2 Respiration}\times n \times M}{m\times t}$$

(3)

where CO_2 Respiration_ is the CO_2_ respiration measurement [ppm], n is the amount of substance, M is the molecular weight of carbon which equals 12, m is the weight of total soil contributing to CO_2_ respiration and t is the incubation time in the closed airtight jar.

In order to calculate the cumulative respiration of the 23 days incubation period, the following equation was used:

$${Cumulative}_{Respiration}= \frac{\sum_{t1}^{t10} {CO}_{2}{-C}_{Equivalent}}{10\times t}$$

(4)

where t1 to t10 are the ten timepoints where respiration measurements were taken [ml g^-1^ h^-^1], divided by 10 to get the average respiration rate per sample, times t which is the total incubation time [h]. In order to make the respiration between the fluctuating and the constant temperature treatment more comparable, a subset of three timepoints (t2, t5, t8) were chosen where all incubation temperatures were at 23°C. The same calculation (4) was used, except that 10 was replaced with 3 because the number of timepoints changed.

The calculations of the microbial carbon use efficiency (CUE) were based on Spohn et al. (2016). The amount of DNA produced during the 48hrs of incubation for each labeled sample is based on the abundance of ^18^O. In a first step the microbial biomass carbon (MBC) was calculated:

${MBC}_{Soil}={DNA}_{Soil}\times11.9$

(5)

Where DNA_Soil_ is the measured DNA concentration [ng ml^-1^] and 11.9 is the conversion factor provided by Spohn et al. (2016).

In a second step MBC produced during the 48hrs of heavy-water inoculation was calculated:

$C_{Growth}= 18MBC= 18O\times\frac{1}{0.31}\times\frac{{MBC}_{Soil}}{{DNA}_{Soil}}\times\frac{1}{m}$

(6)

Where ^18^O [mg] is the oxygen incorporated into microbial DNA during the incubation period and m is the mass of the soil [g]. C_Growth_ was derived from the MBC (eq. 5).

CUE was calculated as follows:

$CUE=\frac{18MBC}{( 18MBC+C_{Respiration})} =\frac{C_{Growth}}{{(C}_{Growth}+ C_{Respiration})}\times100\%$

(7)

where C_Growth_ is the flux of microbial C allocated to biomass production (growth) [ml h^-1^ g^-1^ dry soil] and C_Respiration_ is the flux of C allocated to the production of CO_2_ (respiration) which was converted beforehand to CO_2_-C_Equivalent_ [ml h^-1^ g^-1^ dry soil].

In order to calculate the temperature sensitivity (Q_10_) of CO_2_ respiration, growth and CUE I calculated the Q_10_ value for each sample during the ^18^O-water incubation using the following formula, as suggested by Fang and Moncrieff, (2001):

$Q10=\left( \frac{R_{2}}{R_{1}} \right)^{10/(T2-T1)}$

(8)

Where Q_10_ refers to the change in the response variable with a 10°C change in temperature and R_1_ and R_2_ indicate respiration rates (or growth rates, CUE) separately measured at T_1_ and T_2_. As the temperature was chosen with a 10°C difference, the superposed term in the formula equals 1. In a second step the mean Q_10_ values for the temperatures 13°C-23°C and 23°C-33°C were calculated in order to get just one Q_10_ value per diversity and temperature treatment.

The 95% Confidence Interval (CI) of the Q_10_ value was calculated as follows:

$$95\% CI=\underline{x} \pm z \times\sigma/\sqrt{n}$$

(9)

where $\underline{x}$ refers to the mean of the samples, *z* represents the confidence level 1.96, $\sigma$ is the standard deviation of the population and *n* is the sample size which was 4 or 3 whenever outliers had to be removed beforehand.

**Data analysis**

Statistical analysis were performed in R software version 4.0.2 (R Development Core Team, 2020). Normality of each variable was tested, and log transformed if needed in order to achieve normality of the distribution of residuals and variance homoscedasticity to ensure validity of the test. Outliers were detected and cross-validated with three different methods and only removed if at least two coincided. The methods consisted of a visual inspection of the boxplots, inspection of the residuals vs. leverage plot and if a value was outside of the 1.5x interquartile range for the first and the third quartiles.

To determine the effect of the diversity treatment, temperature regime treatment and the incubation temperature on the C-cycling processes cumulative respiration, DNA yield and 95% CI of the Q10 values of growth, respiration and CUE, a two-way ANOVA was conducted. Whenever the p-value of the ANOVA indicated that at least one of the treatment groups differed from the others (*p* < 0.05) Tukey’s Multiple Comparisons of Means post-hoc test was used to determine which diversity levels were significantly different from each other. Results were considered significant when *p* < 0.05. Within that context, the diversity treatment, the temperature treatments (fluctuating and constant) and the incubation temperature (13°C, 23°C, 33°C) are the independent variables. The dependent variable changed depending on what was looked at. In order to find the best-fit model, the lowest Akaike Information Criterion (AIC) was determined and diagnostic plots were looked at and compared among several models.	In order to quantify the effect of incubation temperature on growth, respiration and CUE a linear-mixed effects model was used. Furthermore t-tests were applied for the two-level factor temperature treatment (constant and fluctuating).
